# Metabolite Interactions Mediate Beneficial Alliances Between *Bacillus* and *Trichoderma* for Effective Fusarium Wilt Control

**DOI:** 10.1101/2025.06.23.660901

**Authors:** Jiyu Xie, Xinli Sun, Tao Wen, Yaoqiang Bai, Tong Qian, Shunjuan Hu, Lihao Chen, Pan Wang, Youzhi Miao, Ruifu Zhang, Ákos T. Kovács, Zhihui Xu, Qirong Shen

**Affiliations:** Jiangsu provincial key lab for solid organic waste utilization, Key lab of organic-based fertilizers of China, Jiangsu collaborative innovation center for solid organic wastes, Educational ministry engineering center of resource-saving fertilizers, Nanjing Agricultural University, Jiangsu provincial key laboratory of coastal saline soil resources utilization and ecological conservation, Nanjing 211800, China; Institute of Biology Leiden, Leiden University, 2333 BE Leiden, The Netherlands

## Abstract

Bacteria-Fungi Interactions play a crucial role in soil nutrient cycling and plant disease suppression. *Bacillus* and *Trichoderma* exhibit antagonism when inoculated on laboratory media, global soil sample analysis reveals a positive correlation between these two genera in addition to enhanced plant-pathogen *Fusarium oxysporum* (FOC) suppression and plant growth promotion. Here, we depict a complexity of interactions in a cross-kingdom consortia of *Bacillus velezensis* and *Trichoderma guizhouense*. Transcriptomic profiling revealed that in the presence of fungi, the key stress sigma factor of *B. velezensis* activates expression of biosynthetic genes for antimicrobial secondary metabolite production. Among these, surfactin induces T22azaphilone production in *T. guizhouense* that hinders oxidative stress. Both surfactin and T22azaphilone contribute to *Bacillus* and *Trichoderma* maintenance in soil in the presence of FOC. Finally, FOC-secreted fusaric acid temporarily inhibits *B. velezensis* growth while it is efficiently degraded by *T. guizhouense*. These metabolite-mediated interactions reveal how competing soil microorganisms could form effective alliances that ultimately enhance plant protection against soil-borne pathogens.

## Introduction

Soil serves as the primary habitat for diverse microbial communities where plant growth-promoting bacteria, beneficial fungi and plant pathogens have coexisted for hundreds of millions of years, leading to complex interactions and metabolite-mediated dialogues. In recent years, Fusarium wilt caused by *Fusarium oxysporum sp.* has emerged as a severe threat to global agriculture, casing devastating losses in economically crops such as cucumber, banana, watermelon and others ^1^. This pathogen not only infects plants and causes wilting through the production of phytotoxins such as fusaric acid ^2^, but also produce chlamydospores to persist in soil for extended periods ^3^, marking it particularly challenging to control. Traditional control methods, particularly chemical fungicides, have shown limited efficacy and led to pathogen resistance and environmental pollution ^4^. Therefore, developing safe and sustainable disease management strategies has become increasingly important.

Building on bacteria-fungi interaction (BFI) insights, biological control has gained attention as an environmentally friendly disease management strategy. Bacteria and fungi have a huge impact on soil ecosystems, and their interactions play a crucial role in regulating microbial communities and maintaining ecosystem functions ^5–8^. These interactions encompass both direct physical contact, such as bacterial colonization on fungal hyphae and biofilm formation ^9,10^, indirect chemical communication through secondary metabolites ^11^ and volatile organic compounds, and competition for nutrients ^12^. Among potential biocontrol agents, *Bacillus velezensis* SQR9, an important plant growth-promoting microorganism (PGPM), can suppress pathogens through various secondary metabolites [12,13] (such as bacillomycin D, fengycin) while promoting plant growth through beneficial goods secretion, phosphate solubilization and beneficial microorganism recruitment ^15^. Similarly, *Trichoderma guizhouense* NJAU 4742 inhibits pathogenic fungi through mycoparasitism, nutrient competition, and hydrolytic enzyme production [15,16], particularly chitinase, while also enhancing plant growth ^18–20^.

Despite the demonstrated potential of these beneficial microorganisms under field conditions ^21^, a molecular understanding of their real-world interaction is lacking. While *B. velezensis* and *T. guizhouense* show strong antagonism against each other in vitro ^10^, these species often coexist stably in natural environments, a paradox that highlights the complexity of soil microbial interactions. The intricate interactions between beneficial and pathogen microbial agents likely involve “alliance formations” among the beneficial microorganisms to maintain the pathogenic fungi at a manageable level. Current BFI research primarily focuses on pairwise interactions rather than how these BFI consortia are influenced by environmental factors and affect other macro- and microorganisms. Only few studies examined so far the multi-interaction mechanisms among plant beneficial fungi and bacteria, and the pathogenic microorganism in soil ^22,23^. These studies have mainly investigated how secondary metabolites influence community assembly and interspecies communication, contributing to the inconsistent efficacy of single-strain applications in field conditions ^24,25^.

Previous studies have demonstrated enhanced plant growth promotion when co-inoculating *Bacillus* and *Trichoderma* species, such as *T. harzianum* and *B. cereus* ^26^, *B. amyloliquefaciens* ACCC11060 and *Trichoderma asperellum* GDFS100 ^27^. However, the underlying mechanisms governing these beneficial interactions remained largely unexplored. Here, we established a unique three-species interaction system comprising *B. velezensis* SQR9, *T. guizhouense* NJAU 4742, and the cucumber pathogen *Fusarium oxysporum f. sp. cucumerinum* (FOC). Using transcriptomic analysis, mutant construction, and chemical analysis by LCMS revealed (1) the role of the SigB sigma factor in *B. velezensis* inducing an arsenal of secondary metabolites; (2) the importance of *B. velezensis* surfactin for enhancement of antioxidant fungal metabolite production in *T. guizhouense*; and (3) the modulatory role of fusaric acid in controlling bacterium versus fungus balance in the tri-species interactions. Our findings not only advance our understanding of microbial interaction mechanisms but also provide scientific basis for developing more effective biocontrol strategies. Additionally, our multi-species interaction research system offers a new paradigm for exploring microbial community assembly mechanisms in complex environments.

## Results

### *Bacillus* and *Trichoderma* positively correlate in soil

To investigate the distribution of the two genera, *Trichoderma* and *Bacillus*, we analyzed 1680 soil metagenomes obtained from various ecological environments (Fig 1A), downloaded from NCBI in February 2025, including agricultural soil (48.9%), rhizosphere (30%), wetland (4.1%), grassland (2%) and unknown soil (15%) (Fig 1B). Through correlation analysis of *Bacillus* and *Trichoderma* abundance, we found a significant positive correlation in overall sample (r = 0.186, p = 1.33e-14, n=1680, Fig 1C). Interestingly, both *Bacillus* and *Trichoderma* showed negative correlations with other agriculturally important microorganisms, including *Aspergillus*, *Glomus*, *Pseudomonas*, *Streptomyces*, *Actinomyces* and *Enterobacter* (Fig S1), highlighting the uniqueness of the positive correlation between these two plant-beneficial microorganisms. Further analysis of each soil environment revealed significant positive correlations across all types, with grassland showing the strongest significance (r = 0.477, p = 0.0043) despite having the smallest sample size (n = 34), followed by agricultural soil (r = 0.251, p = 2.736e-13, n = 821) and rhizosphere (r = 0.166, p = 2e-04, n = 504), while wetland demonstrated the weakest correlation (r = 0.274, p = 0.0228, n = 69).

**Fig. 1.**
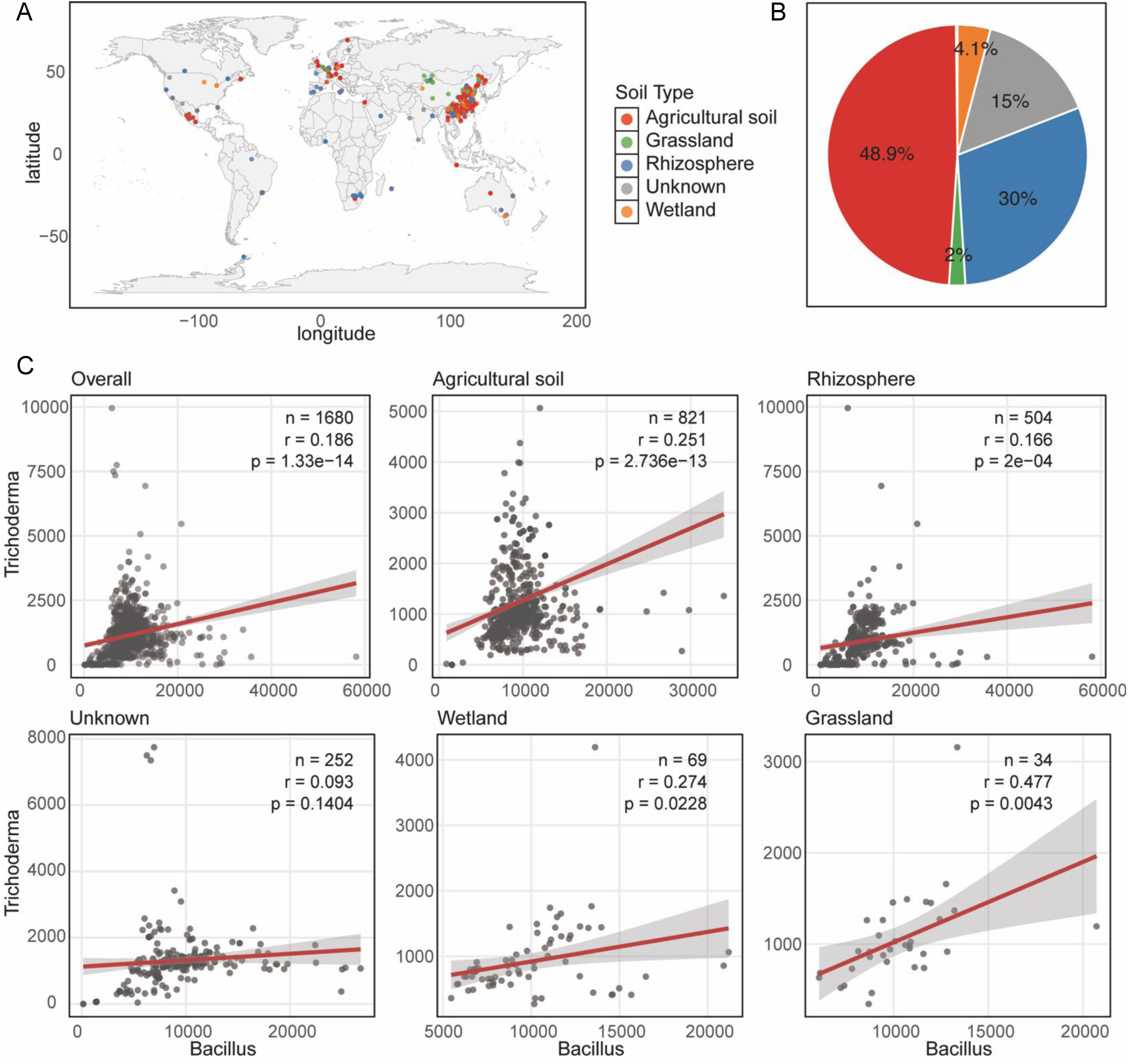
The correlation analysis of *Bacillus* and *Trichoderma* from global soil data. **(A)** The geographic region, soil types used in this study are displayed. **(B)** The ratio of different soil types in the merging data. **(C)** The correlations between *Bacillus* and *Trichoderma* in different soil types. Pearson correlation analysis was performed to assess the correlation between *Bacillus* and *Trichoderma* abundance. Statistical significance was evaluated with *p*-values, where values less than 0.05 were considered statistically significant.

Notably, the strength of positive correlation was higher in agricultural soil than in the rhizosphere, reflecting differences in microbial interaction intensities across different ecological niches. This widespread positive correlation suggested that members of the *Bacillus* and *Trichoderma* genera may have developed a mechanism for mutual adaptation and coexistence through certain interactions, rather than competition.

### Co-inoculation of *B. velezensis* and *T. guizhouens*e enhance disease suppression and plant growth promotion

The interaction among *B. velezensis*, *T. guizhouense* and FOC was first observed on plates (Fig 2A). *B. velezensis* inhibited both fungi, as expected based on the documented production of various secondary metabolites ^28^. In contrast, *T. guizhouense* suppressed FOC growth through mycoparasitism via secretion of chitinase and protease that was previously reported to degrade FOC mycelia ^17^. Given the well-studied capabilities of *B. velezensis* and *T. guizhouense* as PGPM individually ^15,29,30^, we further investigated the effects of their co-inoculation on plant growth in pot experiments.

**Fig. 2.**
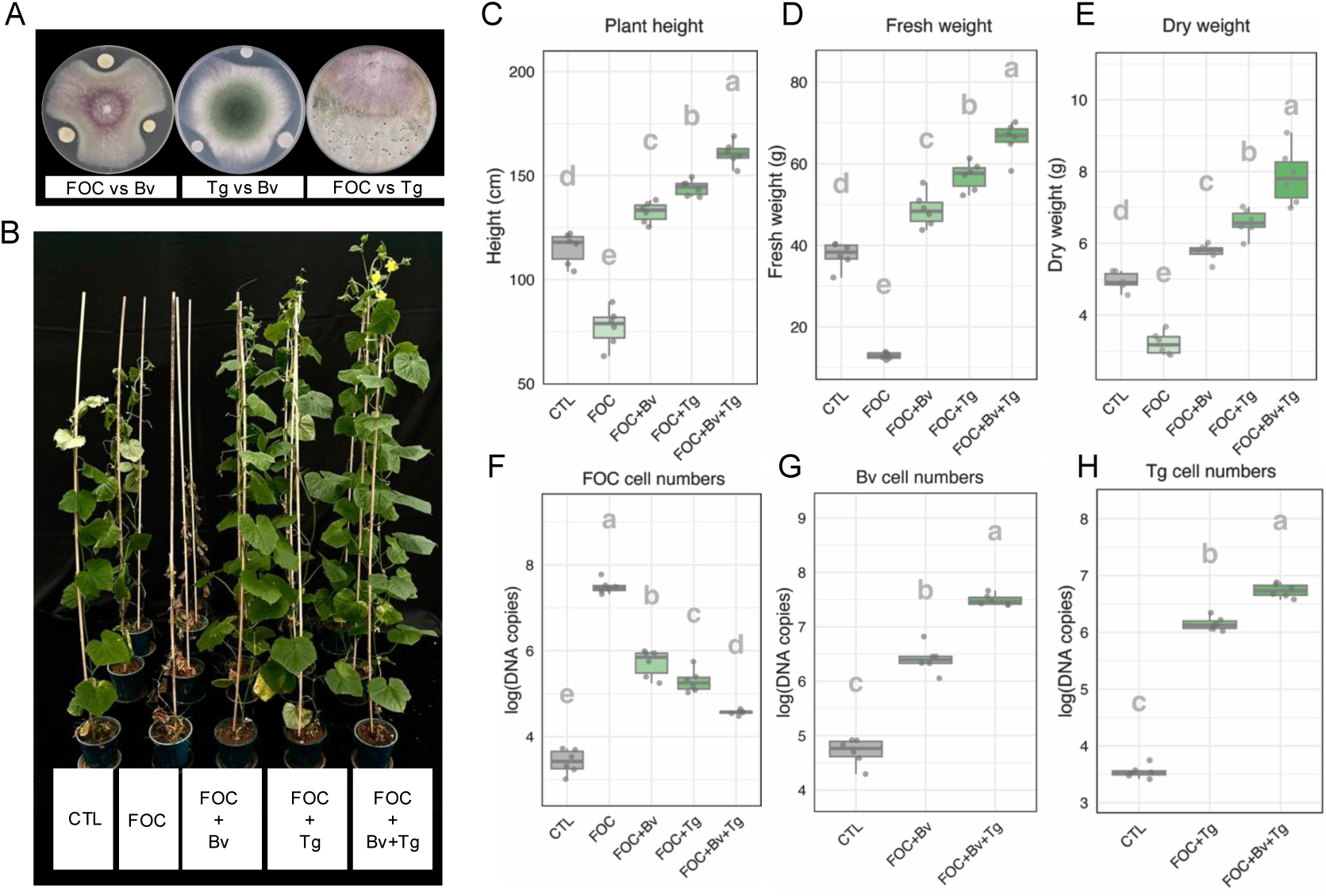
Co-inoculation of *T. guizhouense* and *B. velezensis* inhibited pathogen FOC and improved cucumbers growth. **(A)** the interactions between *T. guizhouense* (Tg), *B. velezensis* (Bv) and FOC. The diameter of Petri dish plate: 9 cm. (**B)** Cucumber plants with different treatments. CTL: control. Plant height **(C)**, fresh weight **(D)** and dry weight **(E)** of cucumber plants. The cell numbers of FOC (F), Bv (G), Tg (H) in soil by RT-qPCR. Bars represent ± s.d. (n=6). Significance test was performed using one-way ANOVA followed by Tukey’s posthoc test. Different letters indicate statistically significant (*p* < 0.05) differences.

We followed control of FOC by different combinations of PGPM and monitored cucumber growth (Fig 2B). After 8-week growth, cucumbers inoculated with FOC alone showed severe yellowing of leaves and growth inhibition, and the plants almost fully withered. Individual inoculation with either *B. velezensis* or *T. guizhouense* not only alleviated FOC-induced damage, but also significantly promoted plant growth compared with the non-inoculated control. Notably, despite the inhibitory interactions between *B. velezensis* and *T. guizhouense* observed on agar medium, their co-inoculation in the presence of FOC (FOC+Bv+Tg) demonstrated the most pronounced disease suppression and plant growth promotion. These synergistic effects were clearly reflected in plant height (Fig 2C), fresh weight (Fig 2D) and dry weight (Fig 2E). We also quantified the microbial populations in the plant soil, which aligned well with plant growth. FOC were lowest in FOC+Bv+Tg treatment (Fig 2F), while Bv and Tg numbers were relatively high in this combination (Fig 2G&H), suggesting potential mutualistic interactions between *Bacillus* and *Trichoderma*, potentially maintaining each other’s abundance, thereby jointly suppressing FOC.

### FOC acts as the primary target in three species interaction

Following the observation of enhanced diseases suppression through co-inoculation of *Bacillus* and *Trichoderma* against FOC, we performed transcriptome analysis to dissect the molecular details of the interaction among these three microorganisms. By sampling the interaction areas between pairwise combinations of these microorganisms (Fig 3A), we scrutinized the distinct transcriptional responses that might be present during their interaction in the plant rhizosphere, although we acknowledge the simplified laboratory setup does not compare to the complexity of soil environment.

**Fig. 3.**
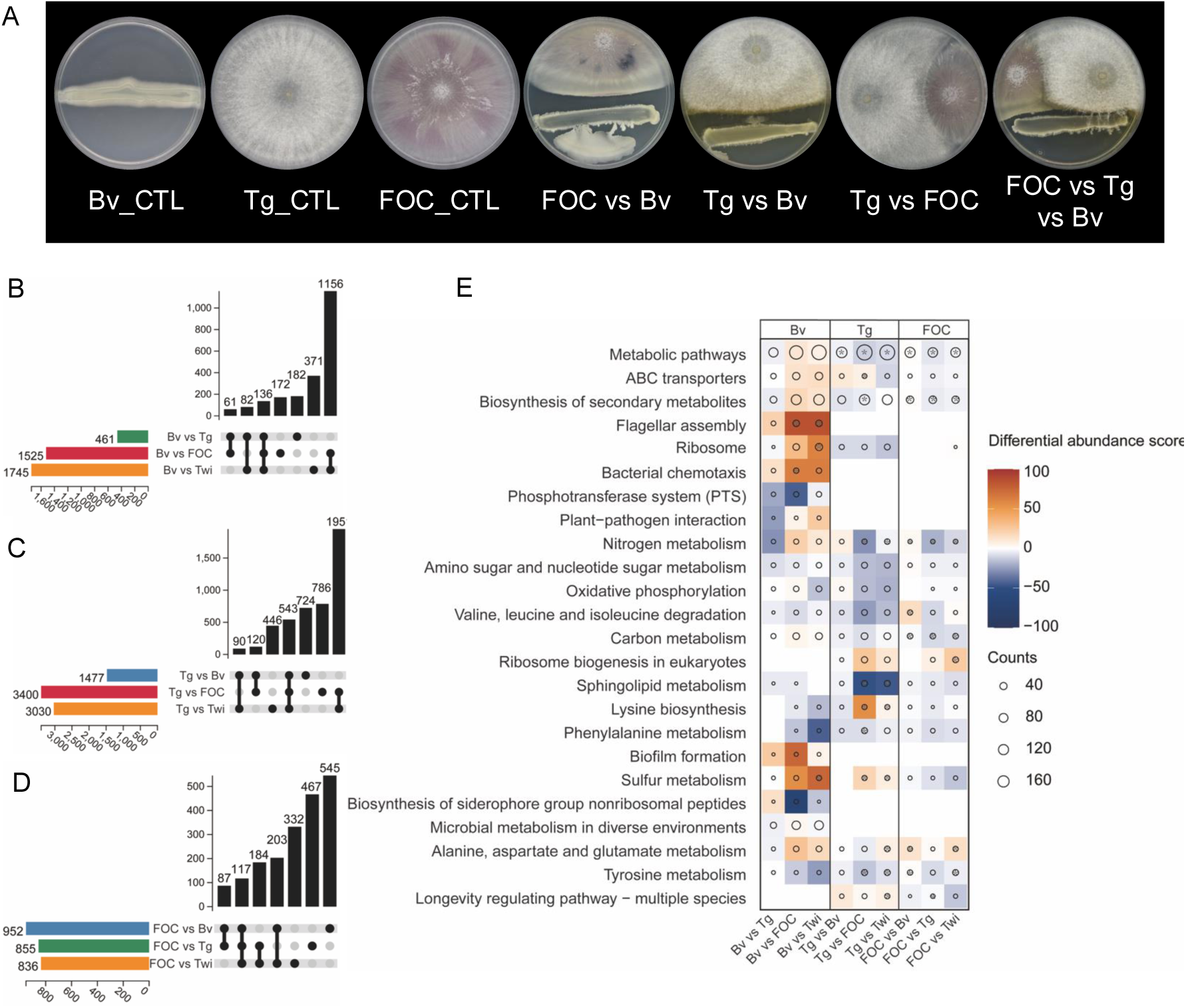
Transcriptome analysis of *B. velezensis*, *T. guizhouense* and FOC. **(A)** Schematic representation of transcriptome treatments. The diameter of Petri dish plate: 9 cm. Numbers of the differentially regulated genes in *B. velezensis* **(B)**, *T. guizhouense* **(C)** and FOC **(D)**. **(E)** KEGG enriched pathway analysis of all treatments in three species. Pathways which LFC>2, FDR< 0.05 were significantly regulated and shown in *.

The differential gene expression analysis showed that both PGPMs exhibited strong transcriptional response against the pathogen FOC. *B. velezensis* exhibited extensive transcriptional changes during interaction with FOC (1525 genes) or in the presence of both fungi (1745 genes), with 1156 genes commonly regulated in both treatments (Fig 3B). Similarly, *T. guizhouense* displayed substantial responses when challenged with FOC (3400 genes) or the presence of both FOC and *Bacillus* (3030 genes), sharing 1951 regulated genes (Fig 3C). Notably, the direct interaction between *B. velezensis* and *T. guizhouense* revealed relatively fewer transcriptional changes (461 and 1477 genes, respectively), supporting the previous finding that despite antagonism on agar medium, these beneficial microbes may effectively coexist. The pathogen FOC showed relatively consistent numbers of differentially expressed genes when interacting with *B. velezensis* (952 genes), *T. guizhouense* (855 genes), or in the presence of both *Bacillus* and *Trichoderma* (836 genes), with distinct gene sets responding to each beneficial microorganism (Fig 3D).

Principal Coordinates Analysis (PCoA) further confirmed that both *B. velezensis* and *T. guizhouense* primarily responded to FOC in the tri-species challenge (Fig S2A&B), consistent with their gene expression patterns (Fig S2D&E). In contrast, FOC exhibited distinct responses to different treatments (Fig S2C&F), indicating its position as the primary target of both PGPMs in the tri-species interaction.

### σ^B^ mediates stress response in *B. velezensis* during interaction with fungi

To deepen our insights into the molecular details underlying the microbial interactions and to understand the functional responses of each species, we performed pathway enrichment analysis (Fig 3E). Building on the transcriptome findings that identified FOC as the primary target, we sought to understand the regulatory mechanisms driving these interactions. Pathway enrichment analysis revealed distinct metabolic and regulatory responses during the interactions of the three species. *B. velezensis* demonstrated unique adaptations through transcriptional upregulations of motility-related pathways, including flagellar assembly and bacterial chemotaxis, particularly when confronting FOC, suggesting enhanced movement ability to detect and respond to environmental signals ^31,32^. Additionally, *B. velezensis* displayed transcriptional response in genes connected to biofilm formation and sulfur metabolism pathways, indicating its competitive strategy through spatial occupation and potential production of sulfur-containing metabolites ^33,34^. *T. guizhouense* exhibited downregulation sphingolipid metabolism and upregulation of lysine biosynthesis, which indicate membrane restructuring and enhanced amino acid metabolism during antagonistic interaction, as reported previously during the interaction between *T. asperellum* YNQJ1002 and *Fusarium graminearum* ^35^. Both *Bacillus* and *Trichoderma* exhibited substantial transcriptional modulation of secondary metabolite biosynthesis pathways, which has been previously reported between *T. asperellum* HG1 and *B. subtilis* Tpb55 in metabolomics experiments ^36^. Basic metabolic processes, including carbon metabolism and amino acid metabolism, were regulated across all three species, with ribosome-related pathways being notably active in both *B. velezensis* and *T. guizhouense*. The common altered transcriptional regulation of ABC transporters encoding genes across all species suggests potential metabolite exchange during these interactions ^37^. Notably, while *B. velezensis* and *T. guizhouense* showed specific and strong pathway regulations, FOC exhibited relatively consistent responses across the different interactions.

Further analysis of functional pathways revealed a transcriptional upregulation of genes associated with σ^B^ (Sigma B). σ^B^ is a global regulator in Gram-positive bacteria primarily responding to environmental and energy stress in various *Bacillus* species ^38^ (Fig 4A). Environmental stress signals are perceived through the stressosome, activating RsbU, which subsequently regulates RsbV and RsbW, ultimately controlling SigB activity ^39^. When *B. velezensis* confronted *T. guizhouense* and FOC individually, the transcript level of *sigB* showed significantly upregulation, potentially triggering rapid stress response. Downregulation of *spo0A* transcript level indicate that *B. velezensis* prioritizes rapid response mechanisms over sporulation during antagonistic interactions ^40^ (Fig 4A). To validate the role of SigB, we constructed a mutant (*ΔsigB*) and an overexpress strain (*OEsigB*). During interactions with both fungi, *ΔsigB* showed a diminished inhibition potency, while *OEsigB* exhibited high antagonistic activity compared with WT (Fig 4B & Fig S3). This was further supported by testing the expression of biosynthetic gene clusters involved in secondary metabolite production in *B. velezensis* (Fig 4C). Collectively, these results demonstrated that SigB is a key regulator mediating rapid stress response in *B. velezensis* during interaction with fungi.

**Fig. 4.**
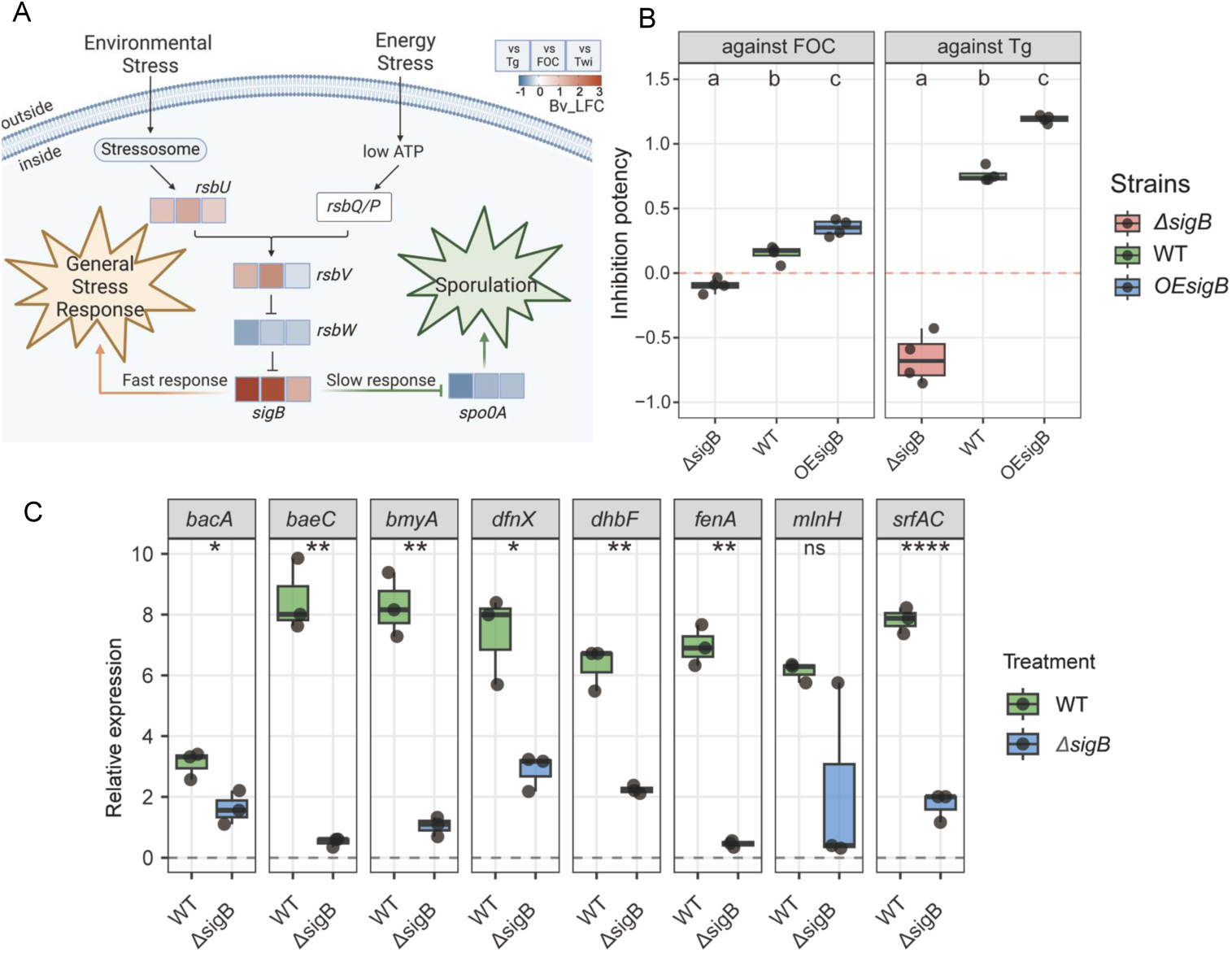
SigB regulated secondary metabolite production to mediate the rapid response of *B. velezensis* to fungi. **(A)** The pathway of global regulator SigB. **(B)** The inhibition potency of *ΔsigB*, WT and *OEsigB*. Bars represent ± s.d. (n=4). Significance test was performed using one-way ANOVA followed by Tukey’s posthoc test. Different letters indicate statistically significant (*p* < 0.05). **(C)** The secondary metabolites genes expression of *B. velezensis* while interacting with fungi, which was quantified by RT-qPCR. Bars represent ± s.d. (n=3). Statistical analysis was performed using *t* test. (\**p* < 0.05, \*\**p* < 0.01, \*\*\*\**p*<0.0001, ns indicates no significant difference). The secondary metabolites of *B. velezensis*: *bacA*: bacilysin; *baeC*: bacillaene; *bmyA*: bacillomycin; *dfnX*: difficidin; *dhbF*: bacillibactin; *fenA*: fengycin; *mlnH*: abolites macrolactin; *srfAC*: surfactin.

### Surfactin enhances T22azaphilone production in *T. guizhouense*

Given that σ^B^-mediated stress response in the presence of fungi activates expression of biosynthetic gene clusters, we tested various *B. velezensis* mutants related to secondary metabolite production (Fig S4). During interaction with *T. guizhouense*, we observed that *B. velezensis* induced the production of T22azaphilone, a yellow compound that has a role in protection of *T. guizhouense* against oxidative stress and fungicides ^41^. In contrast, T22azaphilone production is less pronounced in the presence of the *Δsrf* strain (Fig 5A), as quantified by HPLC (Fig 5B). Surfactin, an important lipopeptide in the *B. velezensis* group species, plays vital roles in collective motility (swarming and sliding) ^42–44^, biofilm formation ^45,46^, interaction with bacteria and fungi ^47–49^, and intraspecies communications ^50,51^. RTq-PCR analysis revealed that expression of the gene involved in T22azaphilone production, *tga5*, was significantly reduced in *T. guizhouense* in the presence of *Δsrf* compared with the wild-type *B. velezensis* (Fig S5). Further experiments demonstrated that *T. guizhouense* produces substantial peroxide (brown) and superoxide (blue) during its interactions with *B. velezensis* and FOC (Fig S6).

**Fig. 5.**
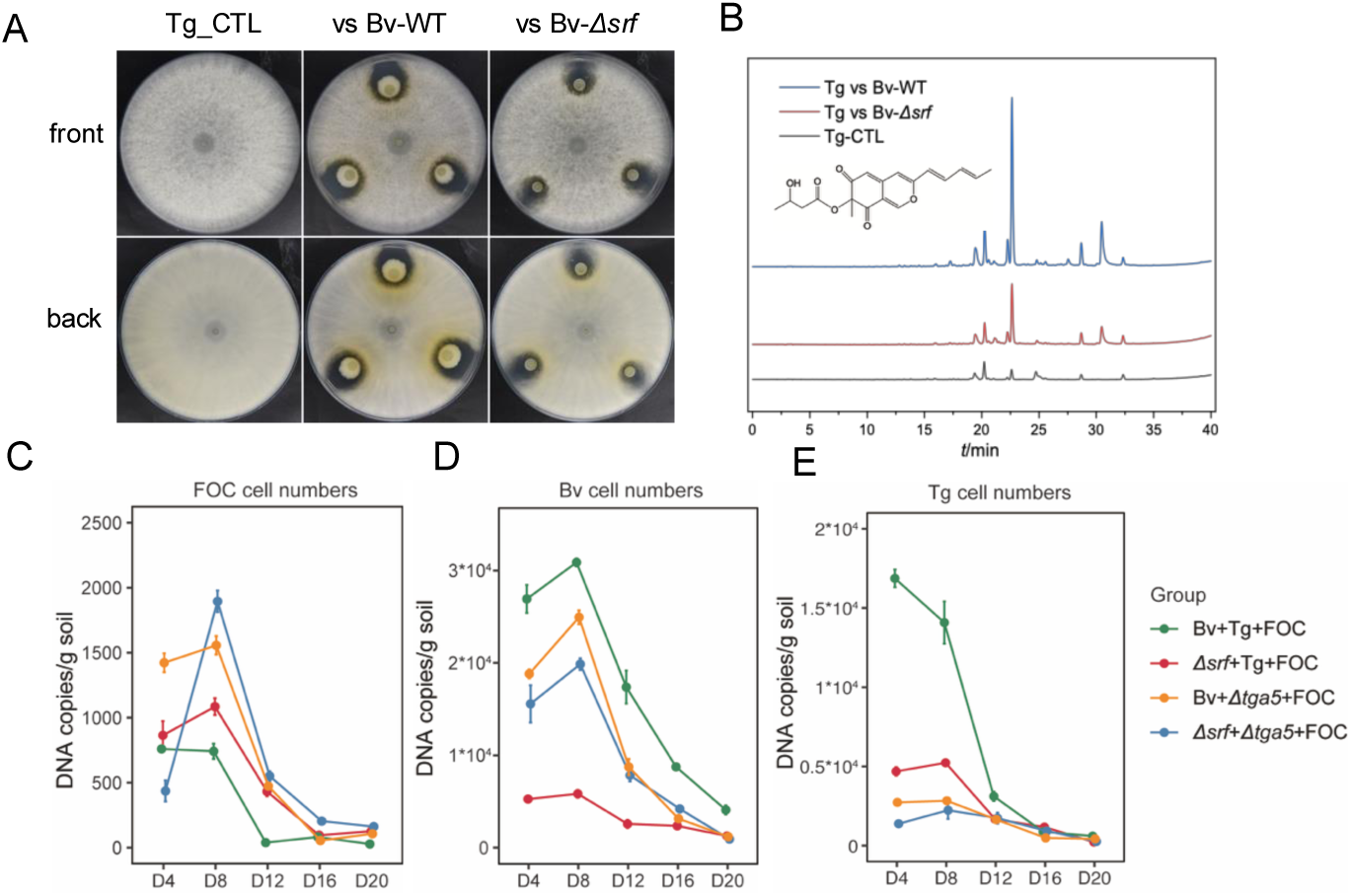
Surfactin induced the protection mechanism of *T. guizhouense*. **(A)** The interactions between *B. velezensis* and *T. guizhouense*. The diameter of Petri dish plate: 9 cm. **(B)** T22azaphilone production quantified by HPLC (peak time= 22.5 min). The FOC **(C)**, Bv **(D)**, and Tg **(E)** cell numbers in sterilized soil experiments with different treatments, quantified by RT-qPCR. Bars represent ± s.d. (n=6).

These in vitro results led us to hypothesize that surfactin might induce a self-protection mechanism in *T. guizhouense* through induction of T22azaphilone production. To validate this hypothesis, we co-inoculated different combination of the three species in soil, including the *Δsrf* and *Δtga5* mutants of *B. velezensis* and *T. guizhouense*, respectively. Over a 28-day cultivation, all three species showed an initial increase of abundance followed by a decline in population sizes. While *B. velezensis* maintained the highest abondance due to its rapid growth compared with the fungi (Fig 5D), *T. guizhouense* populations in the Bv+Tg+FOC were higher than in any other combinations (Fig 5E), supporting our hypothesis: *T. guizhouense* WT maintained higher populations in the presence of *B. velezensis* WT due to a surfactin-mediated induction of T22azaphilone production. In contrast, *T. guizhouense* level was reduced when interacting with *B. velezensis Δsrf*, and displayed the lowest abundance when the *Δtga5* strain was inoculated that lacks T22azaphilone production (Fig 5E). Coincidently, FOC level was lowest in the Bv+Tg+FOC treatment with gradual increase when surfactin or T22azaphilone production was inactivated in *B. velezensis* or *T. guizhouense*, respectively (Fig 5C), due to the suppression in the presence of high abundant wild-type PGPM.

In conclusion, these experiments revealed that bacterial and fungal secondary metabolites enhance abundance of PGPM in soil, potentially enhancing their collective antagonism against the pathogen.

### Fusaric acid-mediated inhibition of *B. velezensis* is prevented by *T. guizhouense*

In addition to the PGPM-mediated influence on the pathogen, we also examined the role of pathogen-derived factors on the interaction of these microorganisms. Fusaric acid (FA), a mycotoxin with low to moderate toxicity against plant and various microorganisms, is the key virulence factor produced by *Fusarium sp.* that plays critical roles in plant pathogenesis ^52–54^. To explore whether FOC can inhibit any of the two PGPM, we investigated the influence of FA. Specifically, to dissect the direct role of FA and prevent the influence of anti-FOC activities of the PGPM on the producing pathogen, we utilized purified FA for our experiments, which displayed corresponding high-performance liquid chromatography peak to FA in the FOC supernatant (Fig S8).

Initial tolerance assays revealed striking differences in FA resistance of *B. velezensis* and *T. guizhouense. T. guizhouense* showed robust tolerance to FA concentrations up to 100 μg mL^-1^ without growth inhibition (Fig 6A). In contrast, *B. velezensis* showed growth delays at FA concentrations exceeding 20 μg mL^-1^, although its growth capacity was maintained below 50 μg mL^-1^ of FA in the first 24 h of cultivation (Fig 6B, S9A). Previous study has reported the potential of *Trichoderma* to degrade FA ^55^. To validated, we co-cultured *T. guizhouense* with 100 μg mL^-1^ FA and quantified FA concentration every 24h using LC-MS (Fig 6C). *T. guizhouense* exhibited efficient FA degradation capacity (Fig 6D). During the first 24h only 10.81% of FA was degraded, while *T. guizhouense* spores germinated at the start of the incubation. However, FA degradation accelerated dramatically to 83.16% by 48h, while FA was almost completely degraded after 72h of culturing (99.76%). This suggested that *T. guizhouense* exerts dual benefit during the plant experiment: in addition to suppressing FOC, it simultaneously eliminates the phytotoxic and anti-*Bacillus* FA.

**Fig. 6.**
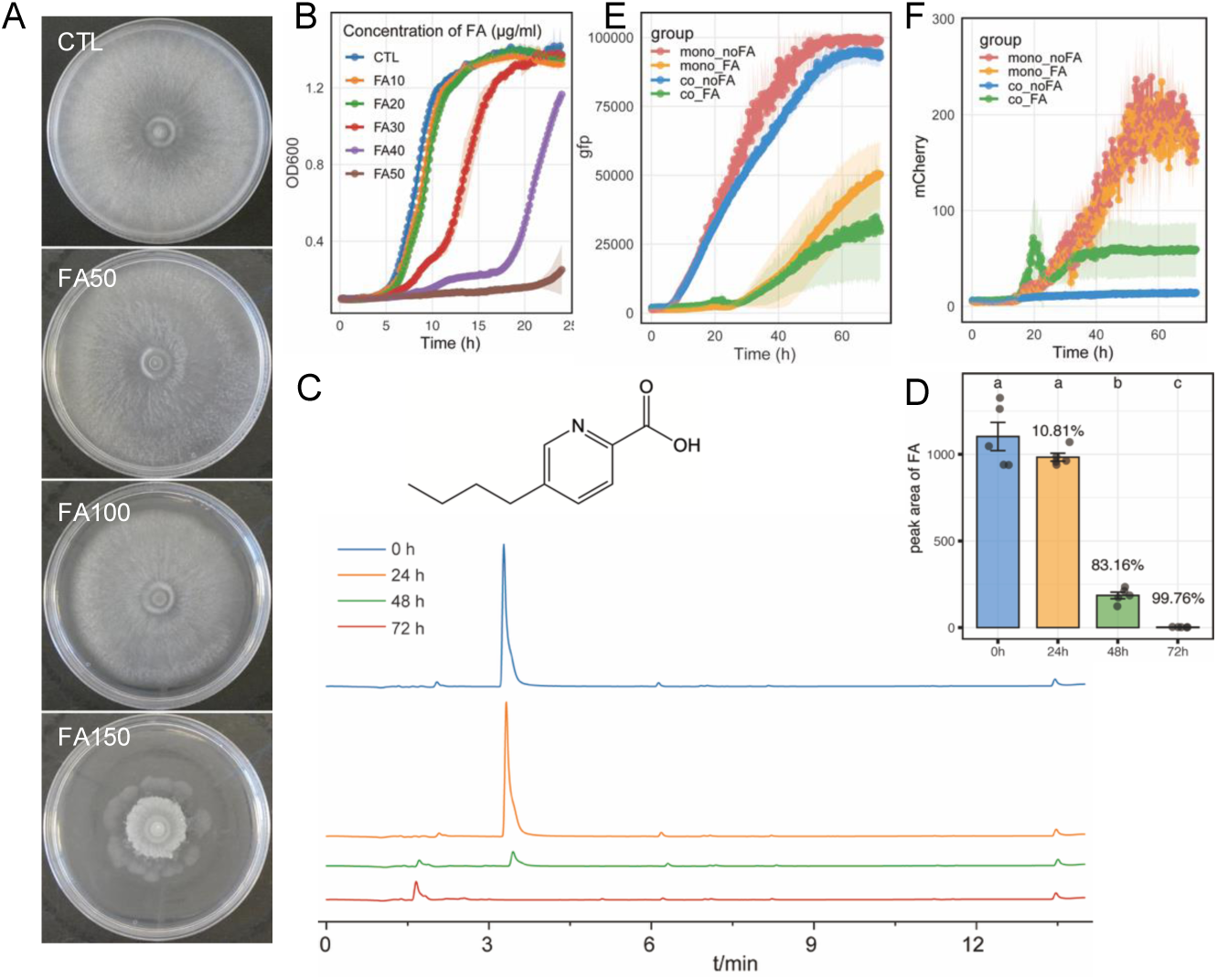
Fusaric acid maintains stability in the tri-species microbial system. **(A)** The growth of *T. guizhouense* in different concentration of fusaric acid. The concentration of fusaric acid: 0, 50, 100, 150 μg mL^-1^. The diameter of Petri dish plate: 9 cm. **(B)** The growth curve of *B. velezensis* in different concentration of fusaric acid. The concentration of fusaric acid: 0, 10, 20, 30, 40, 50 μg mL^-1^. Bars represent ± s.d. (n=3). **(C)** *T. guizhouense* could degrade fusaric acid in 72h, quantified by LCMS (peak time= 3.1 min). **(D)** The peak area of fusaric acid. Bars represent ± s.d. (n=4). Significance test was performed using one-way ANOVA followed by Tukey’s posthoc test. Different letters indicate statistically significant (*p* < 0.05) differences. The growth curve of *B. velezensis* by gfp **(E)** and *T. guizhouense* by mCherry **(F).** Treatment: Mono_noFA, monoculture without fusaric acid. Mono_FA: monoculture with fusaric acid 30 μg mL^-1^. Co_noFA: *B. velezensis* and *T. guizhouense* coculture without fusaric acid. Co_FA: *B. velezensis* and T*. guizhouense* coculture without fusaric acid 30 μg mL^-1^. Bars represent ± s.d. (n=6).

To further follow the interactions among the two PGPM in the presence of FA, we used *B. velezensis* and *T. guizhouense* constitutively expressing GFP and mCherry proteins, respectively. When *B. velezensis* and *T. guizhouense* were cocultured, *T. guizhouense* was unable to growth, while *B. velezensis* growth was unaffected (Fig 6E-F and Fig S9, co_noFA). We attributed this to their different growth rates, as *B. velezensis* reached its stable phase within 24h, severely inhibiting *T. guizhouense* spore germination. However, when 30 μg mL^-1^ FA was supplemented to the coculture, we observed stable coexistence of *B. velezensis* and *T. guizhouense* (Fig 6E-F and Fig S9, co_FA). We hypothesized that FA delayed *B. velezensis* growth (Fig 6B and E), while the *T. guizhouense* spores, unaffected by FA, had sufficient time and nutrients to germinate (Fig 6F). Between 24-48 hours, *T. guizhouense* began degrading FA, gradually alleviating the inhibition of *B. velezensis* by FA, allowing it to resume normal growth.

In conclusion, although FA is typically considered a toxin, it unexpectedly facilitates a stable co-existence between *B. velezensis* and *T. guizhouense* at certain FA concentrations. Once *T. guizhouense* reaches sufficient biomass, it degrades FA that allows collective growth of both beneficial microorganisms, therefore jointly suppressing FOC through various mechanisms, including secondary metabolites and enzymes, and ultimately leading to an effective pathogen suppression and enhanced plant growth.

## Discussion

In this study, we revealed that *B. velezensis* and *T. guizhouense* jointly suppress the plant pathogen FOC in soil, ultimately leading to suppression of pathogenesis and enhanced plant growth. We hypothesize the following interaction among these 3 microorganisms (Fig 7). *B. velezensis* senses the presence of other fungi and initiates a rapid response through σ^B^-mediated enhancement of secondary metabolite production. One of these secondary metabolites, surfactin induce T22azaphilone production in *T. guizhouense* that serves as self-protection and damage reduction. Parallel, fusaric acid secreted by FOC, as a virulence factor, delays *B. velezensis* growth that is reversed upon *T. guizhouense* reaching a sufficient biomass and degrading fusaric acid. Collectively, *T. guizhouense* and *B. velezensis* inhibit the plant pathogenic fungus through hyphal degradation and secondary metabolite-mediated inhibition, respectively, leaving *B. velezensis* and *T. guizhouense* in balance as PGPMs. The combined actions of both beneficial microorganisms effectively suppress the pathogen, establishing a balanced community that promotes plant growth. Our study provides important insights into the complex interactions between beneficial fungi and bacterial in soil, revealing the molecular details towards understanding microbial consortia functioning in agriculture.

**Fig. 7.**
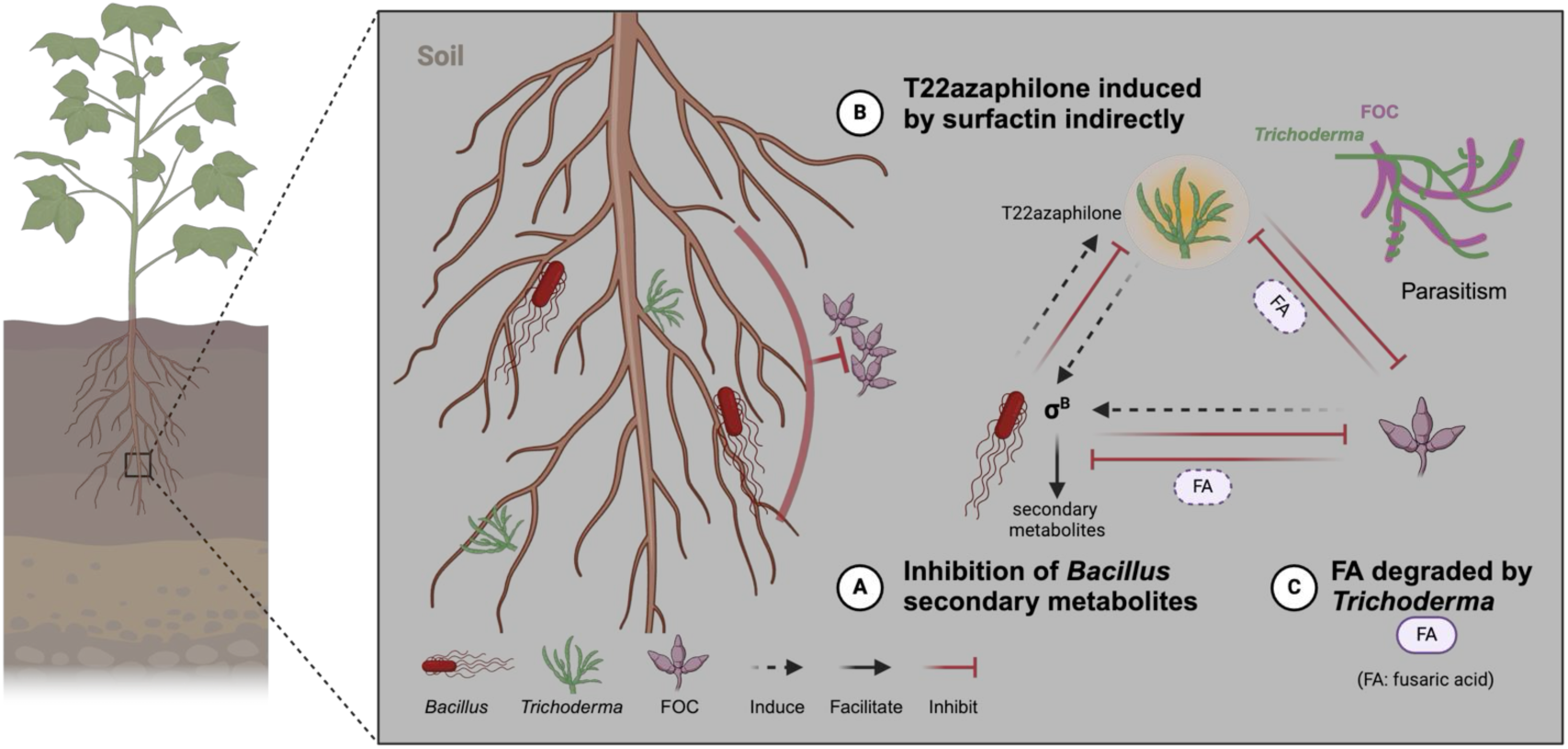
Summary diagram: secondary metabolites mediated molecular interactions and growth. **(A)**The secondary metabolites could inhibit the growth of *T. guizhouense* and FOC. And surfactin induce the T22azaphilone (yellow compound) production, which could protect *T. guizhouense* itself. **(B)** *T. guizhouense* could degrade the hyphae of FOC to inhibit the growth (parasitism). **(C)** Fusaric acid could delay the growth of *B. velezensis* to let the *T. guizhouense* growth first, then degraded by *T. guizhouense*. Plants benefited from reduced pathogen pressure and increased beneficial goods from PGPMs, ultimately enhancing plant growth and health.

Our global soil metagenome analysis revealed significant positive correlations between the *Bacillus* and *Trichoderma* genera across various soil types, providing strong evidence that these beneficial microorganisms naturally co-occur rather than compete in diverse environments. This consistent positive correlation suggests an underlying ecological interaction where these microbial groups potentially engage in complementary activities: *Bacillus* might produce compounds that could affect *Trichoderma* establishment, while *Trichoderma* might modify soil environments in ways that could favor survival of *Bacilli*. The strong positive correlation observed in grassland environments, despite the limited samples, suggested that ecosystems with high organic matter and plant diversity foster more complex and stable microbial networks ^56^. Indeed, *B. subtilis* group species have been shown to have higher abundance in grass land compared with forest soil, paired with a complex co-occurrence network ^57^. The stronger correlation in agricultural soil compared with rhizosphere environments reflects how different ecological niches influence microbial interaction intensities. The observed correlation between *Bacillus* and *Trichoderma* genera across different environments hint a possible synergistic relationship that could be exploited in agricultural systems using co-inoculation strategies, potentially offering more stable biocontrol and plant growth promotion while developing resilient soil microbiomes against pathogen invasion and environmental stresses.

Although *T. guizhouense* and *B. velezensis* exhibit antagonism in vitro, when inoculated at two separate positions on an agar-solidified medium, the transition from antagonism to synergy in the cucumber pot experiments offers a more realistic scenario into their environmental interaction. The increased anti-FOC and plant promotion performance of the co-inoculated strain suggests beneficial interactions between *Bacillus* and *Trichoderma*. Indeed, RNA sequencing highlighted the strongest transcriptional landscape changes in both *B. velezensis* and *T. guizhouense* in the presence of FOC. For example, the σ^B^ global regulator mediated *B. velezensis* rapid response to fungi through upregulating secondary metabolites and chemotaxis while maintaining defensive measures like biofilm formation.

Our study revealed the complex regulation of secondary metabolite production during multi-species interactions. These compounds function beyond simple antagonism, serving as chemical communication signals that mediate ecological relationships. Surfactin, as an important secondary metabolite of *B. velezensis*, has multifunctional role in microbial communities from antimicrobial activity to signaling ^47–49,58^. The concentration dependent effect of these metabolites suggests a fine-tuned balance in natural settings. Particularly noteworthy is T22azaphilone’s dual protective role against both oxidative stress and fungicides ^41^, representing a unique adaptation mechanism compared to other known fungal defense strategies. The induction of protective mechanisms in *T. guizhouense* by surfactin further demonstrates how *Bacillus* metabolites could trigger adaptive responses in *Trichoderma*. Such cross-kingdom chemical communication resembles previous observations where iturin A of *B. subtilis* C2 induces fungal (*Gliocladium roseum* and *T. harzianum*) chlamydospore formation at low concentration, but suppressed the growth of those fungi at high concentration ^59^, supporting the hypothesis that secondary metabolites effects are concentration-dependent, which contributes to maintaining microbial community homeostasis while retaining competitive capabilities when necessary.

Fusaric acid in another example of microbial derived metabolite that has dual role, in addition to toxicity against the host, it also influenced community stability of *Bacillus* and *Trichoderma*. A delayed growth of *B. velezensis* allows *T. guizhouense* germination and growth that subsequently degrades fusaric acid, which exemplifies temporal niche differentiation, a key mechanism allowing species coexistence in natural ecosystems ^60,61^. Similarly, soil bacteria are capable of degrading the toxic oxalic acid produced by *Sclerotinia sclerotiorum* and to reduce its pathogenicity ^8,62^. Our previous research also demonstrated that when *T. guizhouense* is cultured first, *B. velezensis* can form stable biofilms on *Trichoderma* hyphae leading to a co-existence of these two microorganisms ^10^, suggesting a possible mechanism for their continued coexistence that also potentially contribute to FOC suppression.

The temporal dynamics during the interaction of these three species suggests distinct phases of interaction (Fig 7). The early stage is characterized by intense competition and stress responses, evidenced by σ^B^ activation and ROS production. As the community matures, the interaction transition towards a more stable state between *Trichoderm*a and *Bacillus*, including fusaric acid degradation and biofilm formation, respectively. Such dynamic stabilization suggests a fundamental ecological principle where initial antagonistic relationships transition into more balanced associations that maximize collective resilience against environmental perturbations. The dynamic nature of these interactions suggests that sustainable agricultural applications should consider not just which microorganisms to apply, but when and in what sequence to apply them to leverage natural succession pattern. The transition from antagonism to cooperation demonstrates how dynamic microbial relationships can be, challenging simplified views of microbial interactions as fixed and binary, and suggesting that ecological context and temporal dynamics might be equally important as species identity in determining functional outcomes in microbial consortia applications.

Despite the insights acquired from this study on how *Bacillus* and *Trichoderma* interact and function together to suppress plant pathogens, several limitations should be acknowledged. Both *B. velezensis* and *T. guizhouense* produce numerous metabolites beyond the few we examined here (surfactin and T22azaphilone), which may also influence their ecological interactions, despite our data confirms that the compounds we studied play major roles in their interactions. Detailed analysis of transcriptomic data could reveal additional biosynthetic pathways and molecular mechanisms involved in this bacteria-fungi interaction. Natural soil environments contain additional complex microbial communities that likely influence the dynamics between these three microorganisms. Future research should examine how the application of *Bacillus*-*Trichoderma* consortia affects indigenous soil microbiota through microbiome analysis to better understand the broader ecological context of these interactions. Additionally, the molecular mechanisms of surfactin-induced T22azaphilone production should be also dissected, especially how environmental factors modulate these interactions.

In conclusion, our findings suggest a sophisticated chemical dialogue that affects microbial community assembly in the rhizosphere and provides a foundation for designing more effective microbial consortia for sustainable agriculture. Complex ecological communities may self-organize and achieve balance under different environmental conditions. This has important implications for understanding soil ecosystems and community assembly. For biocontrol applications, our insights suggest that successful biocontrol microorganism combinations may require carefully timed species introduction and consideration of metabolic interactions, rather than simply combining organisms with desired traits.

## Materials and Methods

### Strains, growth conditions and interaction on plates

The strains in this study are listed in Table 1. *B. velezensis* (abbreviated below as Bv) SQR9 (China General Microbiology Culture Collection Center, CGMCC No. 5808, NCBI accession No. CP006890.1) was grown at 30 °C in Lysogeny Broth medium (LB-Lennox, Carl Roth, Germany) supplemented with 1.5 % Bacto agar if required. *T. guizhouense* (abbreviated below as Tg) NJAU 4742 (CGMCC No. 12166, NCBI accession No. LVVK01000021.1) and FOC (Agricultural Culture Collection of China, ACCC No. 30220) were grown at 28 ◦C in Potato Dextrose Agar (PDA, BD Difco, US) medium. Spores were harvested with 5 ml water and filtered through Miracloth then stored at 4°C. Media were supplemented with selective antibiotics: chloramphenicol (Cm, 5 μg mL^-1^), zeocin (Zeo, 20 μg mL^-1^), spectinomycin (Spec, 100 μg mL^-1^) and Hygromycin B (Hyg, 100 μg mL^-1^).

**Table 1.** Bacterial and fungal strains used in this study.

| Strains | Genotype | Reference |
| --- | --- | --- |
| *B. velezensis* SQR9 | Wild-type isolate<br>bacterium | 10 |
| *B. velezensis* gfp | wild type with<br>pNW33n-gfp, Cm<sup>R</sup> | 10 |
| *B. velezensis* ΔsigB | ΔsigB::Spec<sup>R</sup> | This study |
| *B. velezensis* OEsigB | pNW33N p43-sigB, Cm<sup>R</sup> | This study |
| *B. velezensis* Δsrf | Δsrf::Zeo<sup>R</sup> | 69 |
| *B. velezensis* Δbac | Δbac::Zeo<sup>R</sup> | 69 |
| *B. velezensis* Δdfn | Δdfn::Zeo<sup>R</sup> | 69 |
| *B. velezensis* ΔfenA | ΔfenA::Zeo<sup>R</sup> | 69 |
| *T. guizhouense* NJAU<br>4742 | Wild-type isolate<br>fungus | 10 |
| *T. guizhouense* mCherry | mCherry, Hyg<sup>R</sup> | 10 |
| *T. guizhouense* Δtga5 | Δtga5::Hyg<sup>R</sup> | 41 |
| *Fusarium oxysporum* f.sp<br>*cucumerinum* | Wild type | 68 |

Cm<sup>R</sup>, Zeo<sup>R</sup>, Spec<sup>R</sup>, and Hyg<sup>R</sup> denote chloramphenicol, zeocin, spectinomycin, and hygromycin B resistance markers.

The interactions among *B. velezensis*, *T. guizhouense* and FOC were analyzed using Potato Dextrose Agar (PDA, BD Difco, USA). To accommodate differential growth rates, initial inoculation consisted of 2 μL fungal spore suspension at the plate center, followed by 24-hour incubation at 28°C. Subsequently, 2 μL of *B. velezensis* overnight culture diluted to an optical density at 600nm (OD^600^) of 1 was introduced. Plates were incubated at 28°C for 3-7 days to monitor interactions. Where applicable, media were supplemented with antibiotics or fusaric acid (Merck, Germany).

### Plant experiments design

Greenhouse trials were conducted from July to September 2023 at Nanjing Agricultural University. Soils for the pot experiments were collected from a historically cucumber-cultivated field with the following properties: pH 6.2, organic matter 26.1 g kg^-1^, available N 171.3 mg kg^-1^, available P 131.4 mg kg^-1^, available K 238.8 mg kg^-1^, total N 1.9 g kg^-1^, total P 1.7 g kg^-1^ and total K 14.2 g kg^-1^, collected from Nanjing, Jiangsu Province, China. Cucumber seeds were surface-disinfected in 2% sodium hypochlorite (3 min), followed by triple rinsing with sterilized distilled water. Seeds were germinated on sterile moistened filter paper in 9 cm Petri dishes at 30 °C. Following one week of germination, seedlings were transplanted into pots containing 2 kg soil and cultivated for an additional week.

The experimental design incorporated five treatments: CTL (control); FOC alone; FOC+Bv; FOC+Tg; FOC+Bv+Tg. FOC inoculation was performed initially, followed by the introduction of *B. velezensis* and *T. guizhouense* after a seven-day interval. Inoculation densities were standardized at 10^7^ spores g^-1^ soil for FOC and *T. guizhouense*, and 10^8^ CFU g^-1^ soil for *B. velezensis*. Plants were maintained under controlled conditions (30°C, 16/8 h light/dark cycles). Plant height, fresh weight, and dry weight were recorded, with 6 replicates per treatment. Cell numbers were quantified by quantitative PCR (qPCR) followed by the methods described below, with 6 replicates per treatment.

### Quantitative analysis of microbial interactions

The inhibitory effects of bacterial metabolites on fungal growth were assessed using a 48-well plate assay system, following previously established methodologies ^63^. The experimental setup involved the preparation of fungal suspensions (*T. guizhouense* and FOC, 10^7^ spores mL^-1^) in Potato Dextrose Broth (PDB, BD Difco, US), supplemented with sterile-filtered (0.22 μm) cell-free supernatant from *B. velezensis* cultures at graduated volumes (0, 10, 20, 40, 60, 80 μL). The cultures were maintained under controlled conditions (28°C, dark) for 72 hours, after which fungal growth was quantified using spectrophotometric analysis. Inhibition potency was expressed through a scoring system derived from the equation: Inhibition Score = 5 - ∑(relative fungal growth).

### Whole-genome transcriptomic analysis and reverse transcription qPCR validation

RNA extraction was performed from interaction zones (Fig 3A) using Trizol reagent kit (Invitrogen, Carlsbad CA, USA), with RNA quality validated using an Agilent 2100 Bioanalyzer. Eukaryotic mRNA was enriched using Oligo (dT) beads, while prokaryotic mRNA was isolated using Ribo-Zero™ Magnetic Kit (Epicentre). Following fragmentation, double-stranded cDNA was synthesized and processed for sequencing. Libraries were sequenced on an Illumina Novaseq6000 platform. The sequencing data were deposited in the NCBI SRA database (BioProject: PRJNA1201065). Following quality trimming, reads were aligned to reference genomes using Bowtie 2-2.2.3 ^64^. Differential gene expression analysis was conducted using DESeq2 R package ^65^, with p-values adjusted for false discovery rate using the Benjamini-Hochberg procedure. Differential expression criteria were established as log2 fold change > 2 and false discovery rate < 0.05. Functional annotation was performed by mapping protein-coding sequences to KEGG Orthology terms via EggNOG-mapper v2 ^66^, with pathway p-values adjusted using Benjamini-Hochberg correction.

For validation studies, RNA samples were reverse-transcribed using PrimeScript RT reagent kit with gDNA eraser (Toyobo). Expression levels of target genes (*recA*, *srfAC*, *bmyA*, *fenA*, *baeC*, *mlnH*, *dfnX*, *bacA*, *dhbF*, *tef*, *tga5*, primers were listed in Table S1 were quantified using ChamQ SYBR qPCR Master Mix (Vazyme). The *recA* and *tef* genes served as internal controls for *B. velezensis* and *T. guizhouense*, respectively. Amplification was performed using an Applied Biosystems Real-Time PCR system. The reaction mixture (20 μL) contained: 7.2 μL H₂O, 10 μL 2× ChamQ SYBR qPCR Master Mix (Vazyme), 0.4 μL of each primer (10 μmol L^-1^), and 2 μL template DNA. Thermal cycling conditions comprised initial denaturation at 95°C for 10 minutes, followed by 40 cycles of 95°C for 30 seconds and 60°C for 45 seconds, with a subsequent melting curve analysis. Relative gene expression was calculated using the 2^−ΔΔCT^ method ^67^.

### Cell number quantification in soil

Sterilized soil (prepared by γ-irradiation (>50kGray, Xiyue Radiation Technology Co.,Ltd., Nanjing, China)) was inoculated with *B. velezensis* (10^8^ CFU g^-1^ soil) and *T. guizhouense*/FOC (10^7^ spores g-1 soil) and at 30 °C in greenhouse. Soil samples were collected at 4-day intervals for DNA extraction using DNeasy Power Soil Pro KIT (Qiagen). Cell numbers were quantified via qPCR by primers listed in Table S1. The single-copy genes were identified through comparative genomic analysis using Roary. Primers targeting these unique genes were designed and validated for specificity by qPCR. The amplified fragments were cloned into pMD19T vectors to generate standard curves showed in Fig S7. Quantitative PCR was performed the same with method above. Each treatment had 6 replicates.

### Construction of Mutant and Overexpression Strains

The *sigB* deletion mutant was generated using an overlap PCR-based strategy as previously described ^68^. The upstream and downstream regions of *sigB* were amplified from wild-type *B. velezensis* genomic DNA by primer pairs sigB_UF/UR and sigB_DF/DR, respectively (Table S1). The spectinomycin resistance cassette was amplified from plasmid pheS-SPC using primers Spc_F/R. These three fragments were subsequently joined by overlap PCR in sequential order (upstream-antibiotic marker-downstream).

For constructing the overexpression strain, the *sigB* gene was amplified using primers OEsigB_F/R and cloned into plasmid pnW33N containing the p43 promoter via EcoRI and BamHI restriction sites. The resulting constructs (deletion PCR products and overexpression plasmids) were directly transformed into *B. velezensis*. Transformants were selected on LB agar supplemented with appropriate antibiotics. All constructed strains were verified by DNA sequencing.

### Growth curve analysis

The growth dynamics of *B. velezensis* were monitored in 96-well plates (Tissue Culture Plate, VWR), with each well containing 200 μL LB medium inoculated with 2 μL overnight culture (OD_600_=1.0). Growth measurements (OD_600_) were recorded at 10-minute intervals over 24/48 hours at 30 °C using an Agilent Synergy H1 microplate reader (Biotek).

For coculture experiments, *B. velezensis* and *T. guizhouens*e were cultivated in 24-well plates (Tissue Culture Plate, VWR) containing 2 mL LB medium per well. Inoculated 1% (v/v) each of B*. velezensis* (OD_600_=1.0) and *T. guizhouense* (10^8^spores/mL). Growth parameters were monitored over 72 hours at 30 °C using scanning mode measurements of OD_600_, GFP fluorescence (excitation/emission: 485/528 nm), and mCherry fluorescence (excitation/emission: 590/635 nm) at 10-minute intervals. Different concentration of fusaric acid was added if required. Each treatment had 4 replicates.

### T22azaphilone quantification

Quantification of T22azaphilone compounds was performed via solvent extraction followed by high-performance liquid chromatography (HPLC). Culture plates were mechanically disrupted, and metabolites were extracted twice with ethyl acetate. The extracts were concentrated to dryness and reconstituted in methanol for chromatographic analysis. HPLC was conducted using a Waters e2695 equipped with an XBridge C18 column (5 μm, 4.6 × 250 mm). Chromatographic conditions were maintained at 30°C with a flow rate of 1 mL min^−1^, employing a linear gradient of acetonitrile/water (containing 0.1% formic acid) from 5% to 95%. Peak time = 22.5 min.

### Fusaric acid degradation experiments

*T. guizhouense* spores (10^7^ spores mL^-1^, 1% v/v) were cultivated in 100 mL minimal medium (MM medium, 10 g L^-1^ glucose, 5 g L^-1^ (NH_4_)_2_SO_4_, 0.6 g L^-1^ MgSO_4_·7H_2_O, 15 g L^-1^ KH_2_PO_4_, 0.05 g L^-1^ CaCl_2_, 0.5 mL L^-1^ trace element stock) at 180 rpm, 28 °C for 72h. FeSO_4_·7H_2_O (0.5 g), MnSO_4_·2H_2_O (0.16 g), ZnSO_4_·7H_2_O (0.14 g), and CoCl_2_·6H_2_O (0.2 g) were dissolved in 50 mL distilled water for the trace element stock. Metabolite extraction was performed by combining 1 mL culture supernatant with 1 mL isopropanol: ethyl-acetate (1:3 v/v) containing 1% formic acid. The extracts were desiccated under N_2_ stream and reconstituted in methanol. Chromatographic separation was achieved using an Agilent Infinity 1290 UHPLC system coupled to an Agilent 6545 QTOF MS with Dual Jet Stream ESI. Samples (1 μL) were separated on a Poroshell 10 Phenyl Hexyl column (250 × 2.1 mm i.d., 2.7 μm; Agilent Technologies) maintained at 40°C. The mobile phase consisted of acetonitrile/water (buffered with 20 mmol L^-1^ formic acid) at a flow rate of 0.35 mL min^−1^, with a linear gradient from 10% to 100% acetonitrile over 15 minutes, followed by a 2-minute hold and 0.1-minute re-equilibration. Data analysis was performed using Agilent MassHunter Qualitative Analysis software.

### NBT and DAB Staining Methods

For superoxide (O^2⁻^) detection, NBT (Nitro Blue Tetrazolium, Aladdin, Shanghai) staining solution was prepared (0.05% NBT in 50 mol L^-1^ phosphate buffer, pH 7.5). For the second method, DAB (Diaminobenzidine, Aladdin, Shanghai) staining solution was prepared (2.5 mmol L^-1^ DAB and horseradish peroxidase (Aladdin, Shanghai)) at 5 purpurogallin units mL^-1^ in 50 mmol L^-1^ phosphate buffer, pH 6.5). For both procedures, 10 mL of respective staining solution was applied to each plate and incubated with gentle agitation (100 rpm) at room temperature for 30 minutes. Following removal of staining solutions, plates were maintained at 25 °C under light protection for 7 hours before imaging. The intensity of blue precipitate in NBT-stained samples indicated O^2⁻^ levels, while brown precipitate formation in DAB-stained samples reflected the oxidative level.

### Metagenomic analysis

The metagenomic data collected from NCBI (field samples’ information was listed in Data S1) underwent quality control analysis with FastQC (v.0.11.9). The data was then processed using Trimmomatic v.0.39 for adapter removal and quality filtering, eliminating bases with quality scores below 15 and sequences shorter than 36 base pairs (with parameters set as: leading: 3, trailing: 3, sliding window: 4:15, minlen: 36). Following quality control, we assembled the cleaned sequences using MEGAHIT, and employed Prodigal (PROkaryotic DynamIc programming Genefinding ALgorithm) to identify open reading frames (ORFs). To generate representative sequences of non-redundant amino acids, we utilized dereps, while Bowtie2 was applied to determine gene abundance. The resulting non-redundant amino acid sequence set underwent annotation for taxonomic classification and functional characterization using multiple databases including KEGG, Kraken2, and GO.

### Statistical Analysis

Analysis and figures were conducted in R 4.4.1, GraphPad Prism 10 and Adobe Illustrator 2024. The significance of differences among multiple groups was determined by one-way ANOVA followed by Tukey’s post-hoc test. The significance between two groups was evaluated using *t* test. For transcriptomic data analysis, differential gene expression was determined using DESeq2 with log₂ fold change > 2 and false discovery rate < 0.05. Correlation analysis between *Bacillus* and *Trichoderma* abundance in metagenomic datasets was performed using Pearson correlation coefficients. Principal coordinates analysis (PCoA) plots were generated using the "vegan" package in R. Sample sizes (n) ranged from 3-6 replicates per treatment as specified in individual figure legends.

## Data availability

Supporting data for all results presented in this paper are contained within the manuscript and supplementary materials. Transcriptome sequencing datasets have been submitted to the NCBI Sequence Read Archive (SRA) database under BioProject PRJNA1201065. Soil metagenomic sequence samples list can be found in Supplementary Data S1. All other data generated and analyzed during this study can be requested form the corresponding author.

## Acknowledgments

This work was financially supported by the National Nature Science Foundation of China (42477310), and the National Key Research and Development Program (2022YFD1500202 and 2022YFF1001800). ÁTK was funded by the European Union (ERC, MicroClock, 101166968). Views and opinions expressed are however those of the author(s) only and do not necessarily reflect those of the European Union or the European Research Council Executive Agency. Neither the European Union nor the granting authority can be held responsible for them.

## Author contributions

J.X., R.Z., Q.S., Á.T.K., Z.X. conceived the project. J.X., X.S., Y.B., T.Q., S.H., P.W., Y.M. performed the experiments. J.X. and T.W. performed Metagenomic analysis. Y.B., T.Q., P.W. performed plant experiments. J.X. and X.S. performed transcriptome analysis. J.X. created bacteria mutants, performed soil-mix and three-species mix experiments. Y.M. performed HPLC analysis. S.H. performed superoxide detection. J.X., Á.T.K., Z.X. wrote the manuscript with corrections from all authors.

