## Supplementary material for "Metabolite Interactions Mediate Beneficial Alliances Between *Bacillus* and *Trichoderma* for Effective Fusarium Wilt Control"

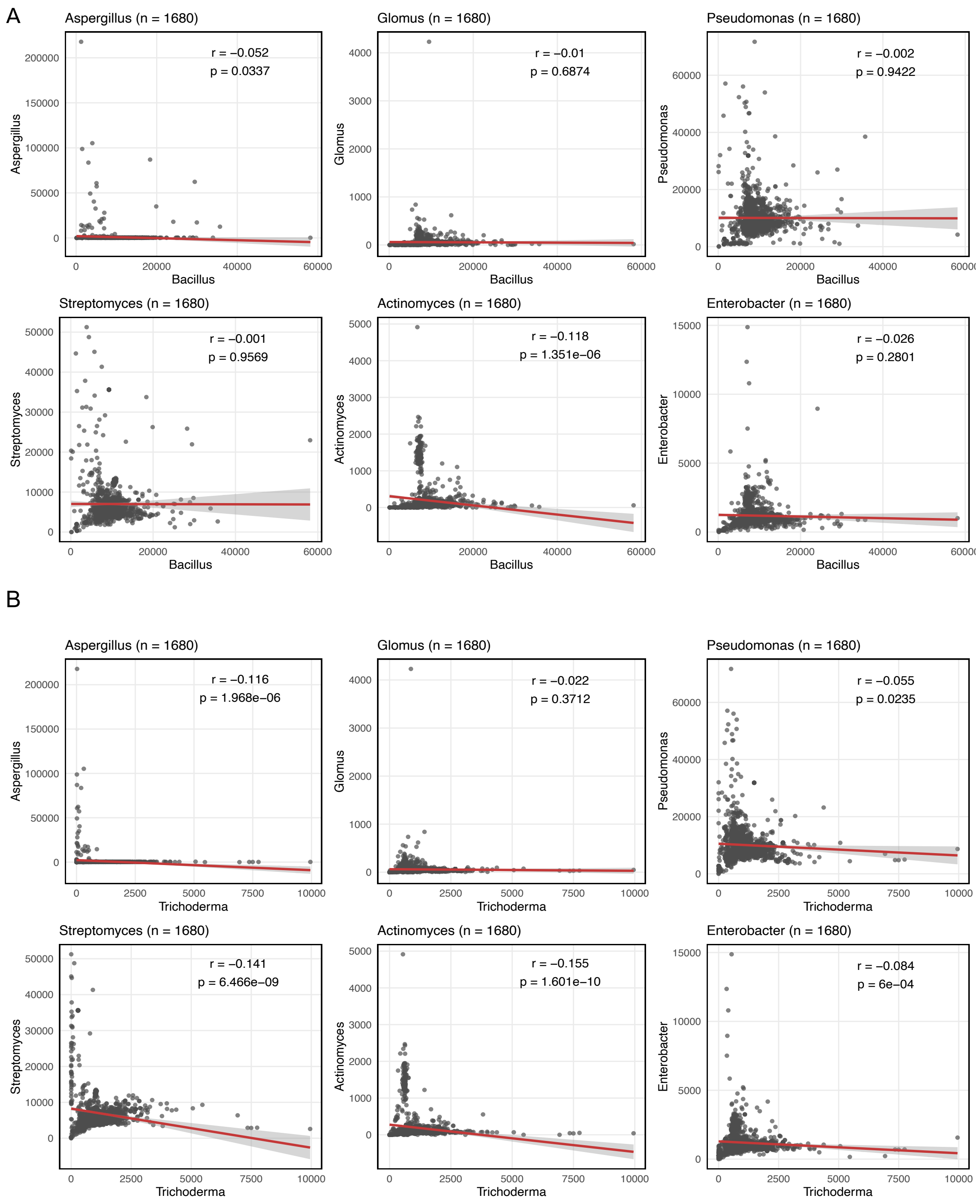


**Fig. S1. The correlation analysis of *Bacillus* (A) and *Trichoderma* (B) with other agriculturally important microorganisms from global soil data.** Pearson correlation analysis was performed to assess the correlation between the abundance of *Bacillus* or *Trichoderma* and other microorganisms (*Aspergillus*, *Glomus*, *Pseudomonas*, *Streptomyces*, *Actinomyces* and *Enterobacter*). Statistical significance was evaluated with *p*-values, where values less than 0.05 were considered statistically significant.

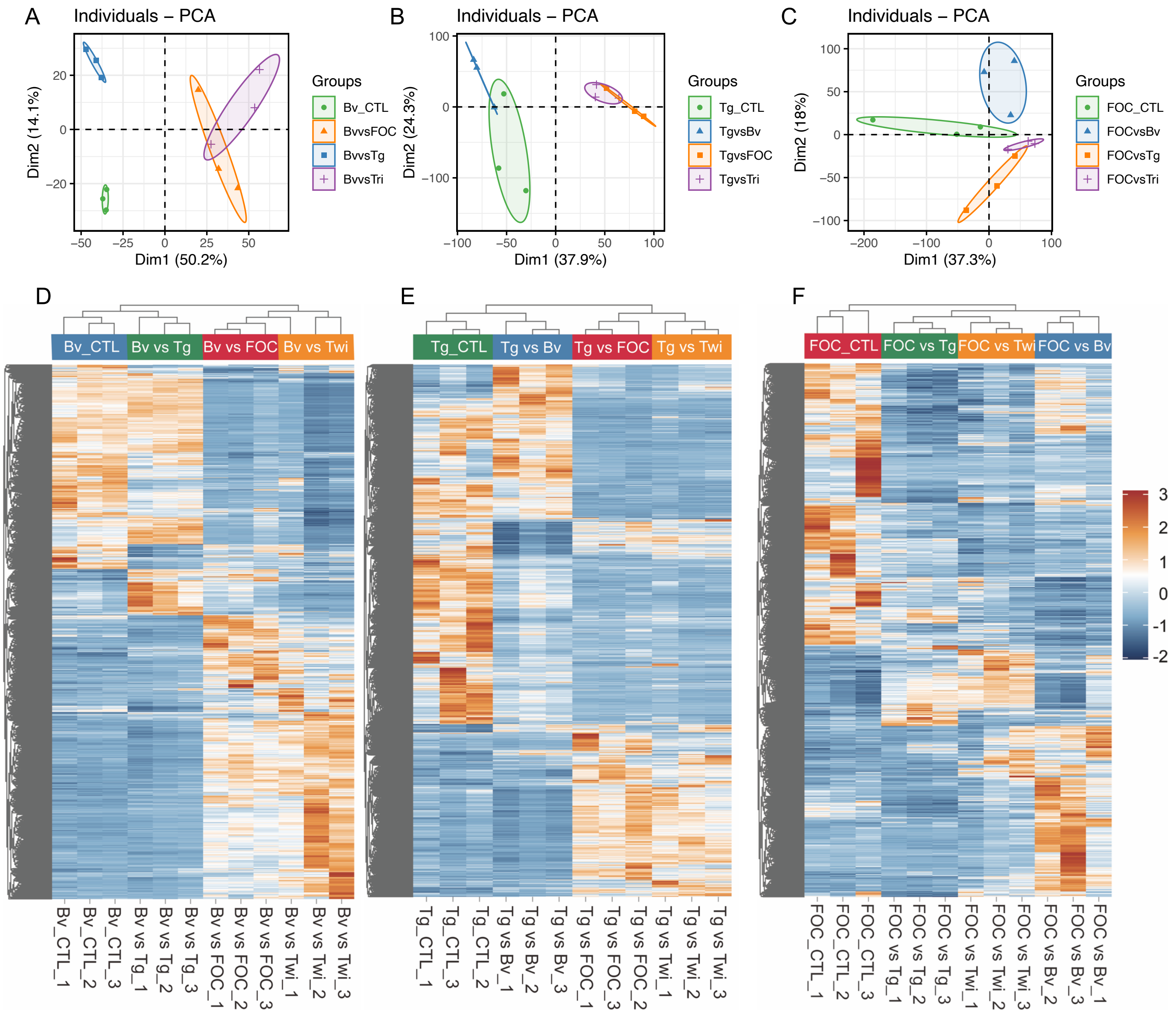


**Fig. S2 Transcriptome analysis.** Principal Coordinates Analysis (PCoA) *of B. velezensis* **(A)**, *T. guizhouense* **(B)** and FOC **(C),** which was plotted based on the Bray Curtis distance metrices for taxonomical data ($p < 0.01$). Permutational multivariate analysis of variance (PERMANOVA) was performed using the adonis function from the vegan R package. Samples were collected from different treatment ($n = 3$). The differential gene expression analysis of *B. velezensis* **(D)**, *T. guizhouense* **(E)** and FOC **(F)** in different treatments.

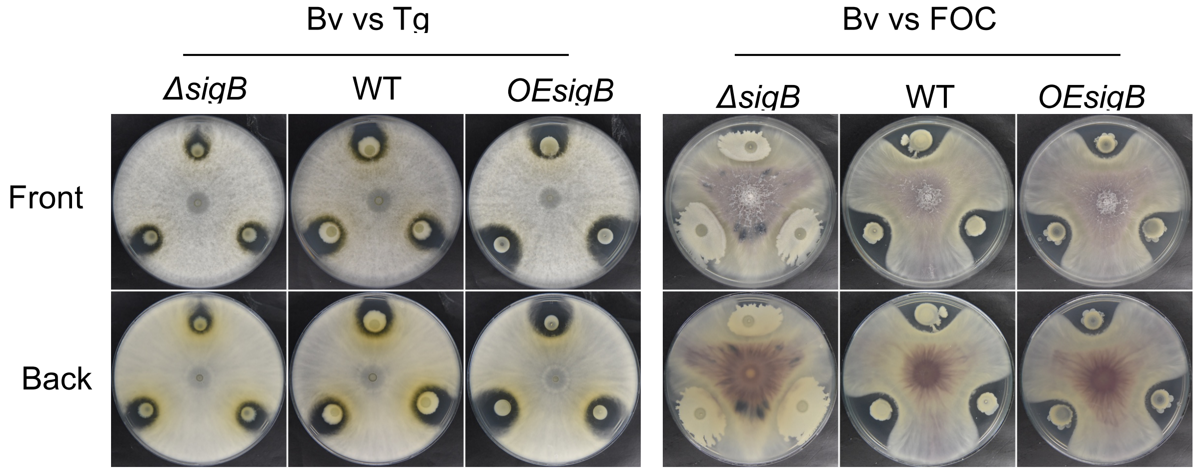


**Fig. S3 The inhibition ability of *ΔsigB* and *OEsigB* to *T. guizhouense* and FOC.** The diameter of Petri dish plate: 9 cm.

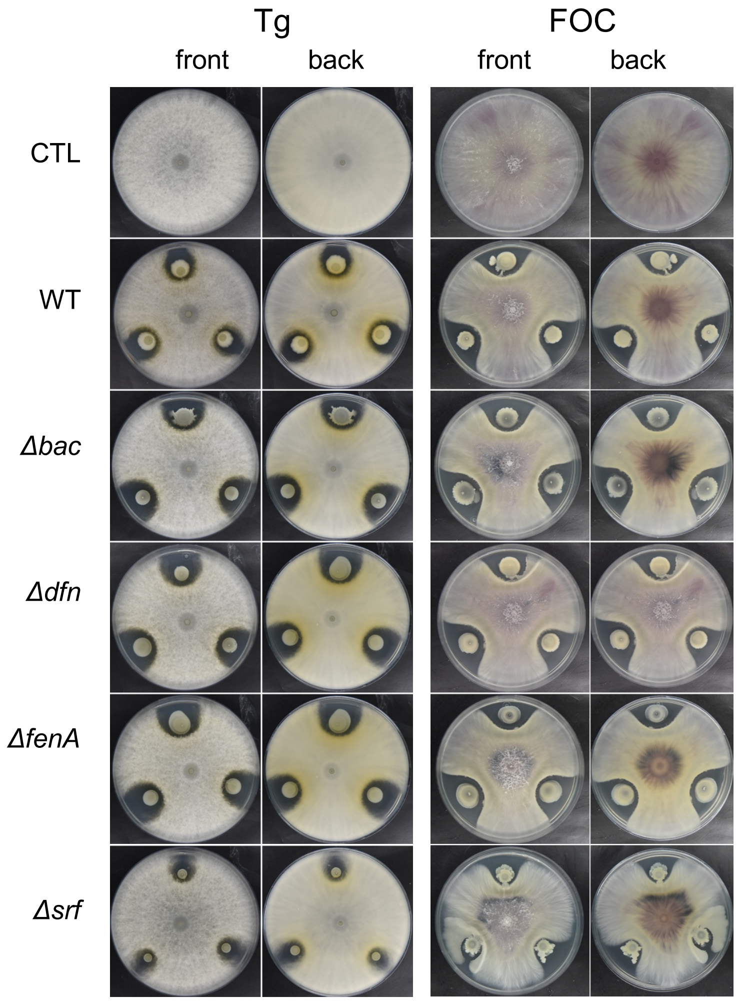


**Fig. S4 The interaction between different B. *velezensis* mutants and two different fungi.** The diameter of Petri dish plate: 9 cm.

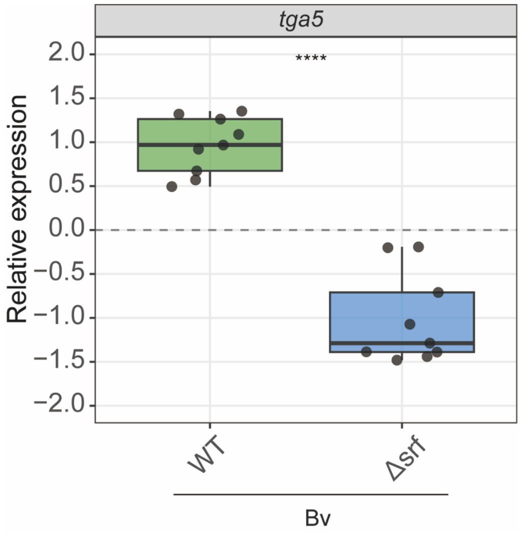


**Fig. S5 The expression of T22azaphilone (*tga5*) while interacting with different *B.velezensis*. Bars represent ± s.d. (n=9).** Statistical analysis was performed using t test (*****p* < 0.00001).

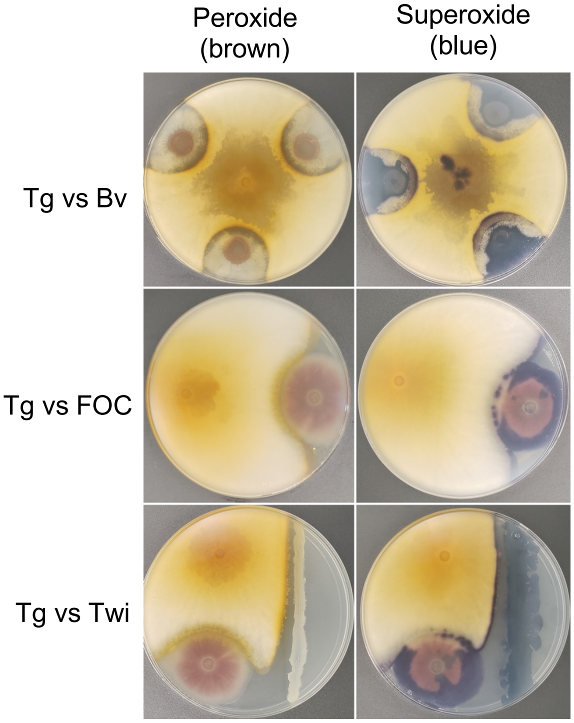


**Fig. S6 The peroxide and superoxide production of T22azaphilone.** Dyeing methods were conducted to verify the production of peroxide and superoxide in *T. guizhouense* while interacting with other species, with staining intensity correlating with substance concentration. The diameter of Petri dish plate: 9 cm.

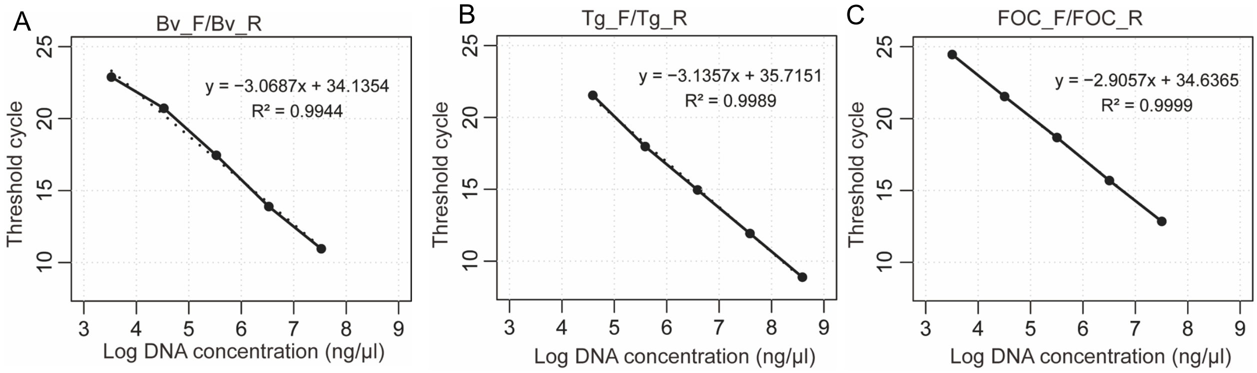


**Fig. S7 The standard curve for quantified cell numbers of *B. velezensis* (A), *T. guizhouense* (B) and FOC(C) by RT-qPCR.**

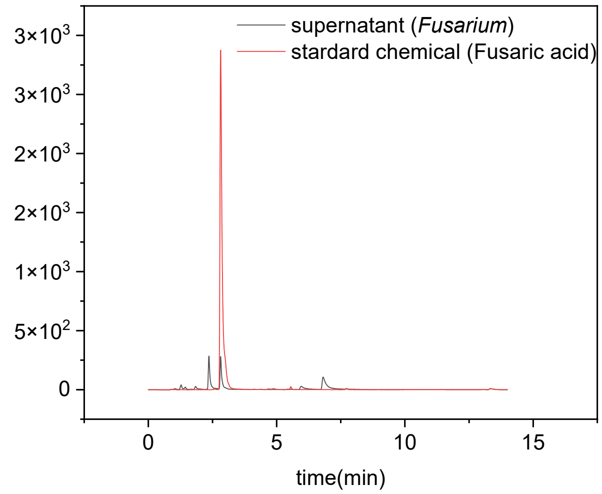


**Fig. S8 Comparison between standard fusaric acid and supernatant by HPLC-MS (peak time: 3.1 min).**

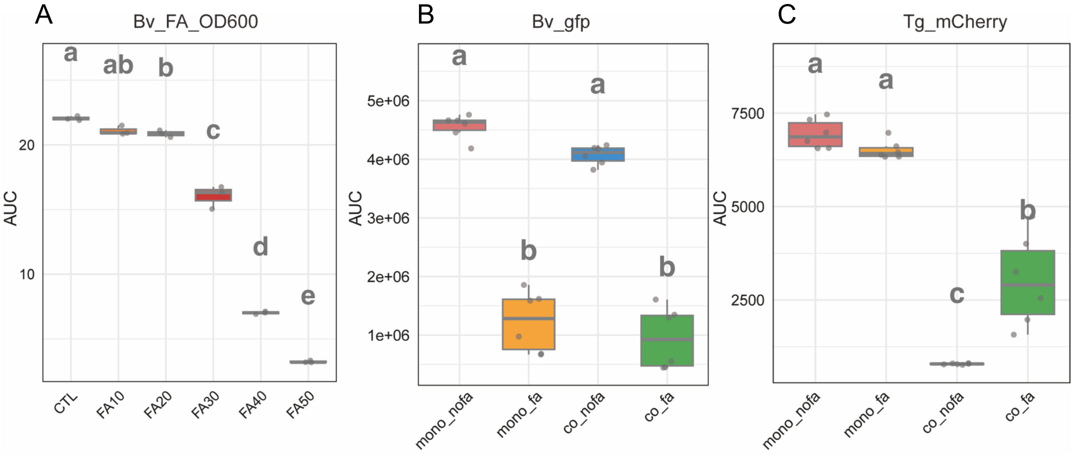


**Fig. S9 AUC (area under curve) analysis of growth curve. (A)** The growth curve of *B. velezensis* in different concentration of fusaric acid. The concentration of fusaric acid: 0, 10, 20, 30, 40, 50 μg $mL^{-1}$. Bars represent ± s.d. (n=3). The growth curve of *B. velezensis* by gfp **(B)** and *T. guizhouense* by mCherry **(C)** in different treatments. Treatments: Mono_noFA, monoculture without fusaric acid. Mono_FA: monoculture with fusaric acid 30 μg $mL^{-1}$. Co_noFA: *B. velezensis* and *T. guizhouense* coculture without fusaric acid. Co_FA: *B. velezensis* and *T. guizhouense* coculture without fusaric acid 30 μg $mL^{-1}$. Bars represent ± s.d. (n=6). Significance test was performed using one-way ANOVA followed by Tukey's posthoc test. Different letters indicate statistically significant ($p$ < 0.05) differences.

**Table S1. Primers used in this study**

| Primers | Sequence (From 5’ to 3’) | Experimental purpose |
|---|---|---|
| **Strains construction** | | |
| sigB_UF | CCGGTGGTTCGTGAAAAGC | To amplify the upstream region of *sigB* gene in *B. velzensis* |
| sigB_UR | CGTTACGTTATTAGTTATCGACG AGTGTTTCCTGGGC | To amplify the upstream region of *sigB* gene in *B. velzensis* |
| Spc_F | GCCCAGGAAACACTCGTCGAT AACTAATAACGTAACG | Cloning |
| Spc_R | GTCCGCATCCGCTTCGATCGTA TAATGTATGCTATA | Cloning |
| sigB_DF | TATAGCATACATTATACGATCGA AGCGGATGCGGAC | To amplify the downstream region of *sigB* gene in *B. velzensis* |
| sigB_DR | GACGGTATTGCCGTCTTCC | To amplify the downstream region of *sigB* gene in *B. velzensis* |
| sigB_F | GCAGCACGGGTAAAAAGCC | To confirm deletion of *sigB* gene in *B. velezensis* |
| sigB_R | CCATTGAGCCGGTCTGCT | To confirm deletion of *sigB* gene in *B. velezensis* |
| OEsigB_F | CGGAATTCATGGCACAACCATC AAAAACTAC (blue site: EcoR I) | To amplify *sigB* gene in *B.velezensis* |
| OEsigB_R | CGCGGATCCTTACATTAGCTCC ATGGAGGGAT (blue site: BamH I) | To amplify *sigB* gene in *B.velezensis* |
| **Relative gene expression quantification** | | |
| RecA_F | AAAAAACAAAGTCGCTCCTCC G | To quantify the relative expression of housekeeping gene *recA* in *B. velezensis* |

| | | |
|---|---|---|
| RecA_R | CGATATCCAGTTCAGTTCCAAG | To quantify the relative expression of housekeeping gene *recA* in *B. velezensis* |
| srfAC_F | CCGCAAACCTTTACTT | To quantify the relative expression of *srfAC* gene in *B. velezensis* |
| srfAC_R | AGGATGTCGGACCAGA | To quantify the relative expression of *srfAC* gene in *B. velezensis* |
| bmyA_F | AGTCTAAGTATTGGCGAAACGA | To quantify the relative expression of *bmyA* gene in *B. velezensis* |
| bmyA_R | ATTATGCTGAAAGTGAAGGGCG | To quantify the relative expression of *bmyA* gene in *B. velezensis* |
| fenA_F | AGCAAGGGAGACACGA | To quantify the relative expression of *fenA* gene in *B. velezensis* |
| fenA_R | CGAGAACCTGGGAGAC | To quantify the relative expression of *fenA* gene in *B. velezensis* |
| baeC_F | CGCACGGATTACATAC | To quantify the relative expression of *baeC* gene in *B. velezensis* |
| baeC_F | AACTCTTGTTTCGCTTC | To quantify the relative expression of *baeC* gene in *B. velezensis* |
| mlnH_F | GAAGGAAAGCGTAGTTG | To quantify the relative expression of *mlnH* gene in *B. velezensis* |
| mlnH_R | GAGGTTCGGAAGATGC | To quantify the relative expression of *mlnH* gene in *B. velezensis* |
| dfnX_F | AGCGGGAGATGAAGTG | To quantify the relative expression of *dfnX* gene in *B. velezensis* |
| dfnX_R | CCGACGGTTGTAATGC | To quantify the relative expression of *dfnX* gene in *B. velezensis* |
| bacA_F | CTGAAGGGACAAGCAGTGAG | To quantify the relative expression of *bacA* gene in *B. velezensis* |
| bacA_R | GATAGGAGACGGGTGGGATA | To quantify the relative expression of *bacA* gene in *B. velezensis* |

| | | |
|---|---|---|
| dhbF_F | AGAGGTTTCGCTATTGG | To quantify the relative expression of *dhbF* gene in *B. velezensis* |
| dhbF_R | TTCGGCTTGTATGTTCC | To quantify the relative expression of *dhbF* gene in *B. velezensis* |
| rsbU_F | TGAACATCAGGTTCCTCCGG | To quantify the relative expression of *rsbU* gene in *B. velezensis* |
| rsbU_R | CCCGTAGCCAATCATGACT | To quantify the relative expression of *rsbU* gene in *B. velezensis* |
| rsbV_F | ATGTATACTCAGCTCCGGTG | To quantify the relative expression of *rsbV* gene in *B. velezensis* |
| rsbV_R | GCCTAAACCCGTACTGTCC | To quantify the relative expression of *rsbV* gene in *B. velezensis* |
| rsbW_F | TCCGGCCGAACCGGAATAT | To quantify the relative expression of *rsbW* gene in *B. velezensis* |
| rsbW_R | CGTTTGTACAAGCTTCGCT | To quantify the relative expression of *rsbW* gene in *B. velezensis* |
| sigB_F | AATGGCGGTGGATCAGCT | To quantify the relative expression of *sigB* gene in *B. velezensis* |
| sigB_R | CGACAGACAGCGCCTGATAG | To quantify the relative expression of *sigB* gene in *B. velezensis* |
| spo0A_F | GAGCTCTGATTCCCGCAGA | To quantify the relative expression of *spo0A* gene in *B. velezensis* |
| spo0A_R | GAGCTCTGATTCCCGCAGA | To quantify the relative expression of *spo0A* gene in *B. velezensis* |
| tef_F | TACAAGATCGGTGGTATTGGAACA | To quantify the relative expression of housekeeping gene *tef* in *T. guizhouense* |
| tef_R | AGCTGCTCGTGGTGCATCTC | To quantify the relative expression of housekeeping gene *tef* in *T. guizhouense* |
| tga5_F | GCTGGATGAGATGCGGAC | To quantify the relative expression of *tga5* gene in *T. guizhouense* |

| | | |
|---|---|---|
| tga5_R | TGTAAGAGGCGGGATGGTG | To quantify the relative expression of *tga5* gene in *T. guizhouense* |
| Cell numbers quantification | | |
| Bv_F | ATCAGGCGTTTAACCGCAAG | To quantify the cell numbers of *B. velezensis* |
| Bv_R | GGCTTCTAGCCTGGCTTTGA | To quantify the cell numbers of *B. velezensis* |
| Tg_F | CTCCACCGGCTGGTGAAAG | To quantify the cell numbers of *T. guizhouense* |
| Tg_R | AAGACCCATCCCAACGCG | To quantify the cell numbers of *T. guizhouense* |
| FOC_F | CATCGCCGTCACCAAAACC | To quantify the cell numbers of FOC |
| FOC_R | GTGCGATCATCCCAATTGGCA | To quantify the cell numbers of FOC |
